# Behavioral and genetic analysis of the effects of the psychedelic 2,5-dimethoxy-4-iodoamphetamine (DOI) in *C. elegans*

**DOI:** 10.1101/2025.03.03.641301

**Authors:** Amanda M. White, Adele D. Bauer, Serge Faumont, Shawn R. Lockery

**Affiliations:** Institute of Neuroscience, 1254 University of Oregon, Eugene OR 97403, USA

## Abstract

Psychedelics show promise in treating depression, PTSD, and substance use disorder, prompting research into their mechanisms of action. Most studies use rodent models, but genetic tools can be challenging to apply. Invertebrate models, like *C. elegans*, offer a cost-effective alternative with short generation times and genetic tractability. This study examined the worm’s response to the psychedelic 2,5-dimethoxy-4-iodoamphetamine (DOI) by assessing four serotonergic behaviors. Effects of DOI exposure on locomotion speed, swimming frequency, and egg-laying were undetectable but DOI strongly inhibited feeding. Interestingly, this effect was independent of serotonin receptors, suggesting DOI may act through alternative pathways. These findings indicate *C. elegans* can serve as a useful model for studying psychedelic drug effects, potentially revealing novel mechanisms beyond the serotonergic system. Further research could help clarify these pathways, improving our understanding of the therapeutic potential of psychedelics and refining their efficacy in treating neuropsychiatric disorders.

## Introduction

Growing preclinical [1–4] and clinical evidence [5–7] suggest that psychedelics are effective in treating conditions such as depression, post-traumatic stress disorder, and substance use disorder [8–10]. A vigorous effort is underway to develop alternative psychedelics with enhanced efficacy [11–13]. At present, this effort is proceeding with a limited understanding of the genetic and biochemical signaling pathways activated by psychedelics. Most of this research utilizes *in vivo* and *in vitro* approaches in rodent models [14]. Pharmacological probes, which activate or antagonize candidate signaling molecules, play a dominant role. Because of selectivity concerns regarding pharmacological probes, genetic approaches have also proven to be valuable, especially gene knockouts. However, deploying genetic tools in rodents can be challenging because of high costs, long generation times, and the relative difficulty of epistasis analysis (using double mutants to order genes into pathways). A complementary approach is to use genetically tractable invertebrate models, which are inexpensive, have short generation times, and are amenable to epistasis analysis. Several key genetic and biochemical signaling pathways were first discovered in *Drosophila melanogaster* and *C. elegans* before being recognized as fundamental to human biology including the Notch [15], programmed cell death [16], RNA interference [17], and Hedgehog pathways [18]. A small number of studies have examined the effects of psychedelics on behavior in *Drosophila* and *C. elegans* [19–22]. *C. elegans* is an advantageous model for studying the effects of psychedelics on the nervous system. *C. elegans* has a compact nervous system of only 302 neurons, which has been fully mapped. Its neurons utilize essentially all of the signaling molecules forming the neurochemical backbone of the mammalian brain [23]: the neurotransmitters acetylcholine, GABA, glutamate, serotonin (5-HT), dopamine, octopamine (similar to norepinephrine [24]), and many neuropeptides (including homologs of oxytocin, vasopressin, and opioids [25,26]). Classic psychedelics are serotonin receptor agonists. *C. elegans* has six serotonin receptor genes which encode orthologs of five of the seven serotonin receptor gene families in humans. Four of the six *C. elegans* receptors are G protein-coupled receptors (*nematode*/HUMAN: *ser-1*/HTR2, *ser-4*/HTR1 & HTR5, *ser-5*/HTR6, *ser-7*/HTR7). The remaining two receptors (*mod-1*, *lgc-50*) are ligand-gated ion channels genetically unrelated to HTR3, the sole ligand-gated serotonin receptor in mammals.

The first step in developing a new model for studying the mode of action of psychedelics is to identify behavioral phenotypes induced by the drug. In this study, we examined the effect of the phenylalkylamine psychedelic 2,5-dimethoxy-4-iodoamphetamine (DOI), a serotonin receptor agonist, on crawling, swimming, egg-laying, and pharyngeal pumping. Toxicity testing showed that DOI was non-toxic up to 3.0 mM. Effects of DOI exposure on locomotion speed, swimming rate, and number of eggs laid per worm were not detectable under our experimental conditions. In contrast, DOI strongly inhibited pharyngeal pumping in a dose-dependent manner that was independent of serotonin receptors. We conclude that DOI can modulate behaviors in *C. elegans* and may act on pathways that do not include serotonin receptors.

## Materials and Methods

### Cultivation and Strains

Nematodes were grown at 20 °C on nematode growth medium plates (NGM; 51.3 mM NaCl, 1.7% agar, 0.25% peptone, 1 mM CaCl_2_, 12.9 μM cholesterol, 1 mM MgSO_4_, 25 mM KPO_4_, pH 6) seeded with 200 μL of a saturated *E. coli* (OP50) liquid culture. All experiments were performed on young adult hermaphrodites synchronized by allowing 10-20 gravid hermaphrodites to lay eggs for six to eight hours. Worms were washed and transferred in M9 buffer (3 g KH_2_PO_4_, 6 g Na_2_HPO_4_, 5 g NaCl, 1 ml 1 M MgSO_4_, H_2_O to 1 liter). Strains used in this study are given in Table 1.

**Table 1.**
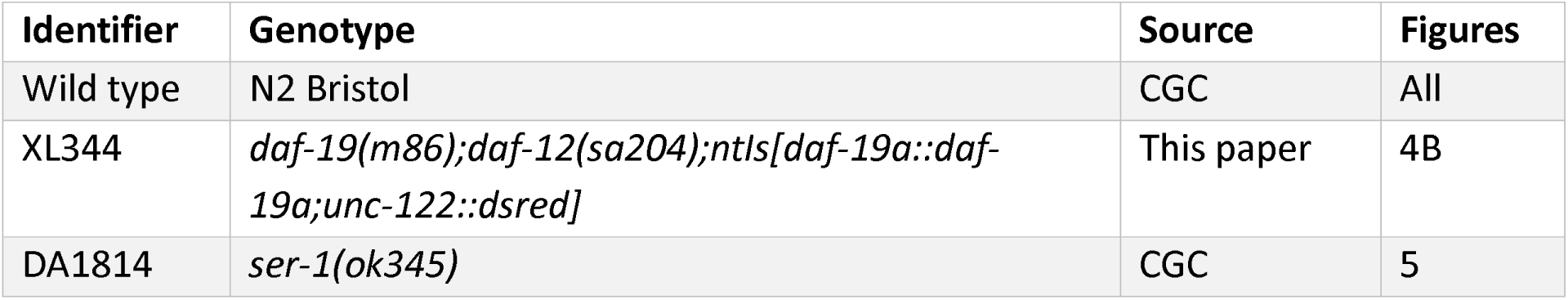
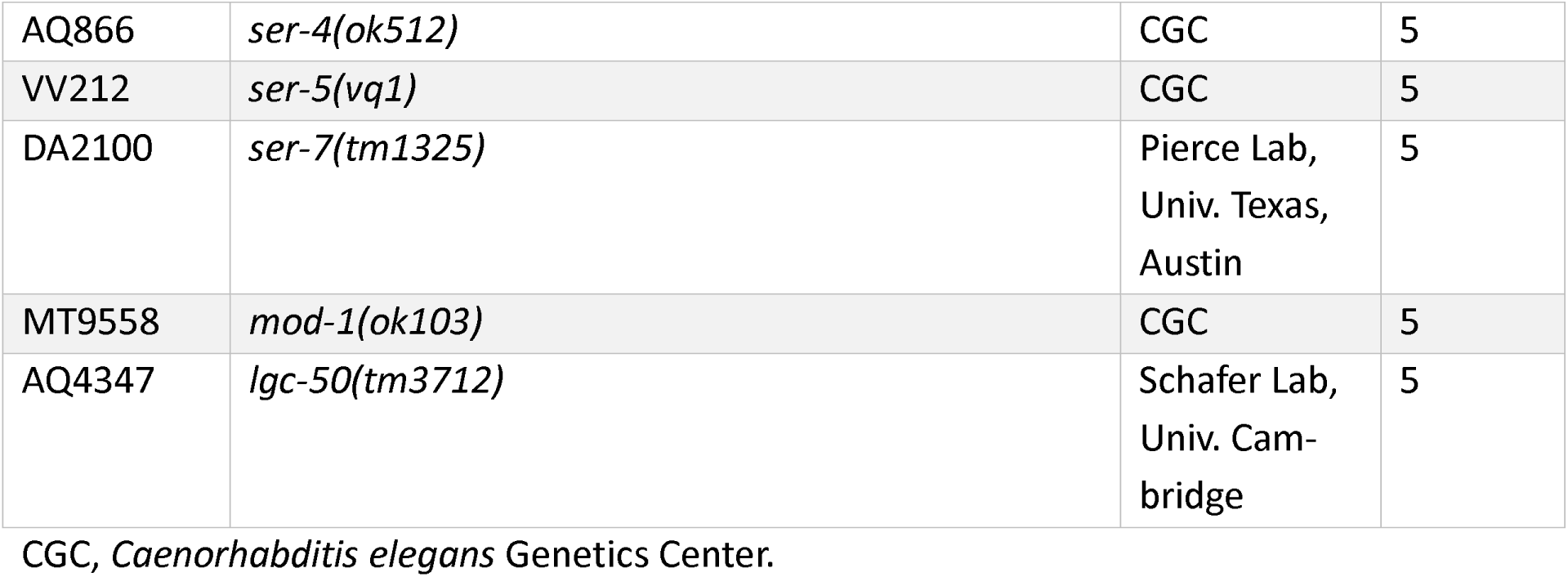
Strains.

### DOI Exposure

2,5-dimethoxy-4-iodoamphetamine (DOI) hydrochloride (Sigma-Aldrich, D101) solutions were prepared by dissolving DOI in M9. For consistency across toxicity, crawling, and egg-laying assays, the same procedure was used: 10–15 young adult worms were transferred by picking from a culture plate to a 2 mL conical glass tube containing 50 µL of DOI (0.1, 0.3, 1, 3, or 10 mM) or vehicle (M9 buffer) solution for 30 minutes before behavioral testing. The collection of worms in each tube comprise a *cohort*. Exposure in the swimming and pharyngeal pumping assays necessitated different procedures which are described below.

### Toxicity Assay

Following exposure, 10–15 worms were transferred by glass capillary pipette to an NGM plate with food. Viability was assessed after 24 hours. Worms were considered to have survived if they displayed spontaneous locomotion and/or exhibited reversals or head bends in response to gentle nose tap with a platinum wire pick.

### Crawling Assay

Following exposure, approximately 10-15 worms were transferred by glass capillary pipette to the inside of a 14 mm (I.D.) copper ring placed on an unseeded NGM plate. After 5 minutes of acclimation, worms were imaged at 23.4 μm/pixel using a macro lens (AF Micro-Nikkor 60 mm f/2.8D, Nikon, Japan) and behavior was video recorded for 10 minutes using a microscope camera (HDMI 1080P HD212, AmScope, Irvine, CA) at 30 fps. Vehicle controls and drug-exposed groups were tested in interleaved fashion. Videos were downsampled to 5 frames/second for analysis. Individual worms were tracked using WormLab (MBF Bioscience, Williston, VT). Tracks were smoothed by Kalman filtering. Average speed values for each track were computed using custom code written in Igor Pro (Wavemetrics, Lake Oswego, OR).

### Swimming assay

A 7 X 4 well array was constructed by punching 3 mm holes in a clear plastic sheet (50 um thickness). The plastic sheet was affixed to a glass plate. Worms were transferred to a blank plate and left to crawl away to remove transferred food. They were then transferred to a well (one worm/well) in 2.5 μl M9 and a 1 min. baseline video was recorded. Then, a 2.5 μl drop of M9 was added to half of the wells to serve as vehicle controls, and a 2.5 μl drop of DOI (2 mM in M9 leading to a final concentration of 1mM) was added to the other half of the wells. The well plate was covered with a glass coverslip to reduce evaporation. A second 1 min. video was recorded within 1-2 minutes (*t* = 0) of adding DOI. Finally, a third 1 min. video was recorded 30 min. after the first one. To simplify the analysis, all videos were background subtracted. To accomplish this, a background image was constructed by averaging all the video frames and that image was subtracted from all frames in th.e video. Behavior was analyzed in Wormlab (MBF Bioscience, Williston, Vermont, USA). All tracks were visually inspected and manually corrected to yield a single continuous track for each worm. The wave initiation rate (number of body waves, i.e., body bends, initiated from either the head or tail/min.) was computed for each worm [27].

### Egg-Laying Assay

Following exposure, the solution containing worms and eggs was transferred by glass capillary pipette to an unseeded NGM plate. Worms and eggs were counted on a stereomicroscope after the solution was fully absorbed into the medium. Vehicle controls and drug-exposed groups were tested in parallel.

### Pharyngeal Pumping Assay

Voltage transients associated with pharyngeal pumping were recorded using a ScreenChip microfluidic system (InVivo Biosystems, Eugene, OR, USA) as described previously [28,29]. Worms were incubated for 30 minutes in an M9 suspension of bacterial food (OP50; OD_600_ 2.5) that contained 1 mM DOI or vehicle only. After incubation, worms were loaded into the reservoir of a microfluidic device that was filled with the incubation solution. Individual worms were then sequentially transferred from the reservoir to the recording channel. Each worm was acclimated to the channel for one to two minutes before pharyngeal pumping was recorded for five minutes. All recordings were performed within 90 minutes after loading animals into the device. Vehicle controls and drug-exposed groups were tested in interleaved fashion. Mean pumping frequency was extracted using custom code written in Igor Pro (Wavemetrics, Lake Oswego, OR, USA).

### Statistical Analysis

G*Power Version 3.1.9.6 [30] was used to determine the minimum sample size for each experiment, using pilot experiments as a guide. Statistical analyses were performed in R version 4.4.2 [31]. One-way ANOVAs were used to assess the effect of drug concentration on toxicity, locomotion, and egg-laying. As pumping frequencies were not normally distributed, significance in feeding assays was assessed via nonparametric tests. Mann-Whitney tests, corrected for multiple comparisons via the Bonferroni method, were used to assess the effect of drug concentration on pumping frequency. Brunner-Munzel tests (implemented in the R package RankFD) were used to evaluate the role of serotonin receptor mutations in the effect of DOI on pumping frequency. Brunner-Munzel tests [32], Bonferroni-corrected for multiple comparisons, were also used as post-hoc tests to evaluate the effects of individual mutations. Outliers were not removed. The threshold for significance was 0.05. In box-and-whisker plots, the box represents 1st and 3rd quartiles, the notch represents the median, the dot represents the mean, and the whiskers indicate the range of data. Details of the statistical analyses, including mode of replication, number of replicates, means, test statistics, *p*-values, effect sizes, are given in S1 Table. For experiments in which there was no detectable effect of DOI, we computed the minimum detectable differences between means, which are given in S2 Table.

### Conversion of Dosage Units

To convert dosage in mg/kg to μM, we used the formula

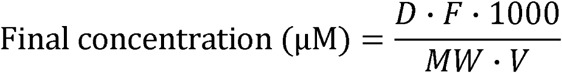

where *D* is the dosage (mg/kg), *F* is the unitless bioavailability of an intraperitoneal injection (0.75), *MW* is the molecular weight of DOI (321 g/mol), and *V* is the volume of distribution in a mouse (≈ 0.7 L/kg).

## Results

### DOI Is Non-Toxic at Low to Moderate Concentrations

We assessed DOI toxicity using a worm survival assay. The worm’s body is encased in a thick, collagenous cuticle that acts as a significant diffusion barrier [33,34]. Accordingly, drugs are often applied at millimolar concentrations. Young adult worms were exposed to DOI in M9 buffer (see Materials and Methods) in the absence of food at various concentrations (in mM: 0.1, 0.3, 1.0, 3.0, 10.0) for 30 minutes; control worms were exposed to vehicle only. After exposure, worms were removed from the solution, placed on a standard culture plate with food, then examined for survival 24 hours later. Spontaneously crawling worms or those that responded to a gentle head touch were considered viable. We found a significant overall effect of DOI concentration on survival (Fig 1; S1 Table, row 1). Post hoc testing showed that survival was indistinguishable from the control group over the range of 0.1 to 3.0 mM; however, at 10.0 mM, survival was significantly reduced (S1 Table, row 2). We conclude that DOI is not overtly toxic at concentrations < 3.0 mM.

**Fig 1.**
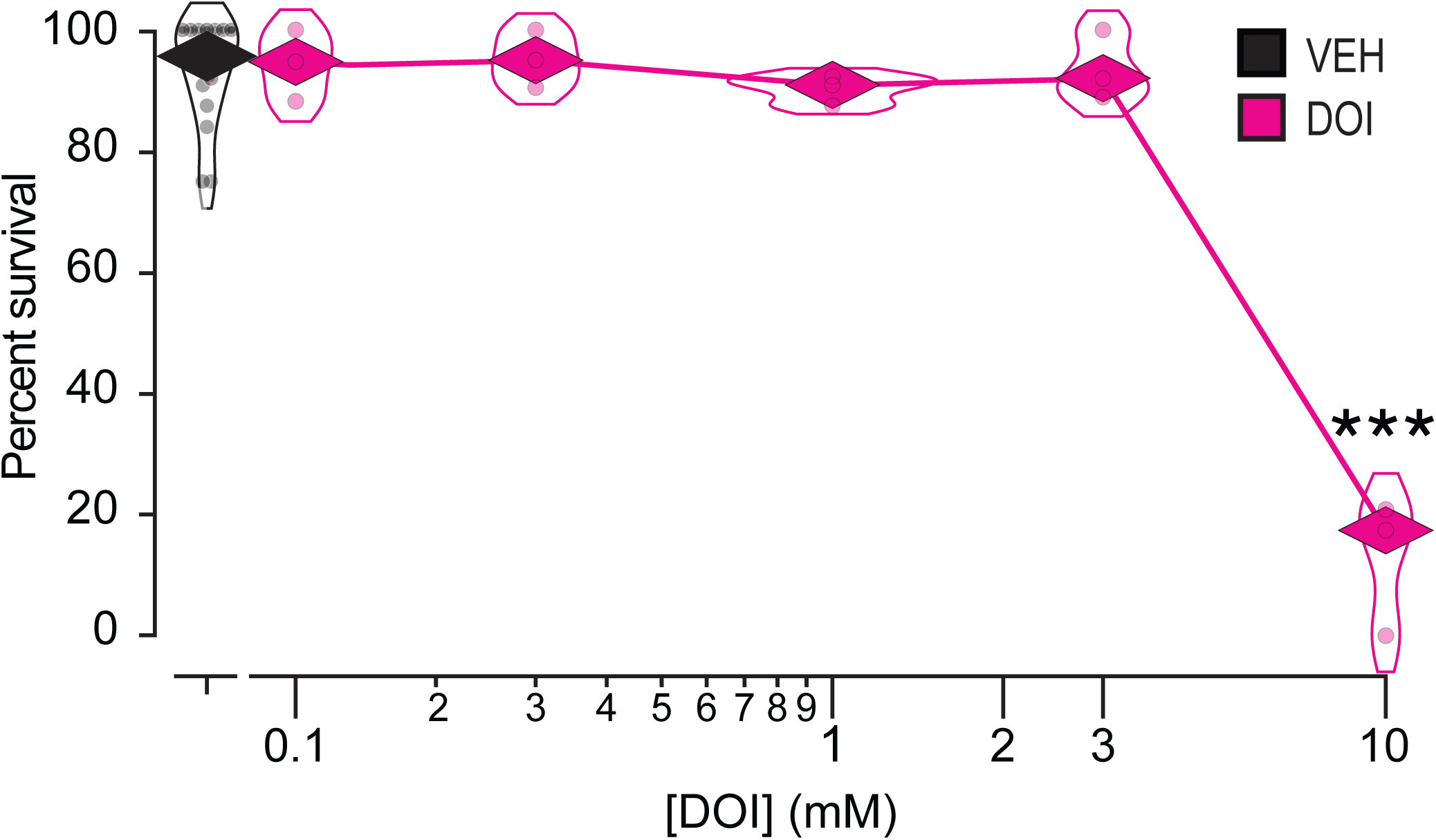
Worm-survival assay. Mean percent survival 24 hours after a 30-minute exposure to DOI. Percent survival is plotted against DOI concentration. Each data point represents the mean of a cohort of 10-15 worms. *n* = 3 cohorts for DOI and 15 for VEH (vehicle). There is a significant main effect of DOI dose. See S1 Table, row 1. *Diamonds*, means; ***, Dunnett’s test, significantly different from VEH, *p* = 8.0 × 10^-7^. See S1 Table, row 2.

### Absence of DOI Effects on Locomotion

*C. elegans* has two main modes of locomotion, crawling and swimming. To test whether DOI affects crawling, young adult worms were exposed to M9 solutions containing DOI at various concentrations within the non-toxic range (in mM: 0.1, 0.3, 1.0, 3.0) for 30 minutes; control worms were exposed to vehicle only. After exposure, worms were tracked as they crawled on foodless culture plates. Behavior was quantified in terms of crawling speed (absolute value of instantaneous speed). Statistical analysis revealed no effect of dosage on mean crawling speed. Statistical analysis revealed a significant effect of time but no effect of dosage and no time × dosage interaction (S1 Table, row 4). Such a decline in crawling speed has been reported previously [35]. Thus, our ability to detect the decline in speed over time indicates that the assay was functional; it can reveal systematic changes in dependent variables. We conclude that DOI has no detectable effect on crawling speed under our conditions.

The crawling assay has two potential drawbacks: (i) worms are tested in the absence of external DOI, potentially reducing its effect and (ii) control and drug-treated animals represent different populations of worms which increases the variance in the data. We therefore devised a swimming assay in which each worm is immersed in the drug solution throughout the experiment and serves as its own control. This was done by placing each worm in a small well (3 mm in diameter) containing 2.5 uL of M9 without drug and video recording baseline swimming behavior for 1 minute. Next, worms in the drug group received 2.5 uL of M9 containing DOI to a final concentration of 1 mM in the well; worms in the control group received an equal volume of M9 only. One or two minutes later, behavior was recorded a second time for 1 minute, then again 30 minutes after the baseline recording. Behavior was quantified in terms of the frequency of initiation of wave-like body bends that comprise *C. elegans* swimming behavior. Statistical analysis revealed no effect of DOI on wave initiation rate (Fig 2B; S1 Table, row 5). We conclude that DOI has no detectable effect on swimming rate under our conditions. Together, the results of the crawling and swimming assays make it unlikely that DOI affects locomotion as we measure it.

**Fig 2.**
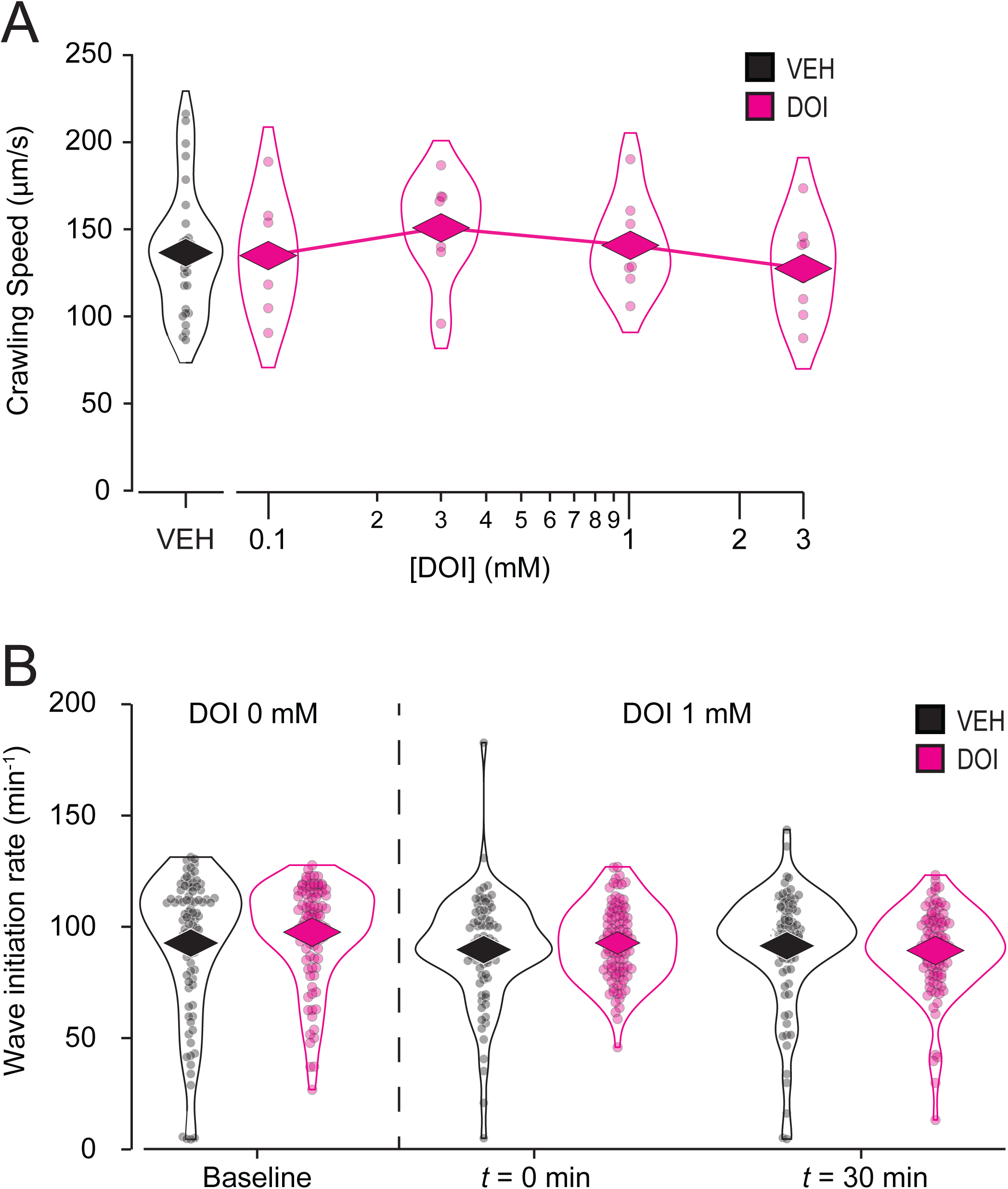

### Absence of DOI Effects on Egg laying

We then tested whether DOI affected egg-laying. We exposed young adult worms to M9 solutions containing DOI at various concentrations in the non-toxic range (in mM: 0.1, 0.3, 1.0, 3.0) for 30 minutes; control worms were exposed to vehicle only. We then pipetted the solution onto a foodless culture plate and counted the number of eggs laid during exposure as well as the number of worms to compute the number of eggs laid per worm. A positive control using serotonin at 28 mM (5mg/ml), a dose known to stimulate egg-laying [36], showed a significant increase in number of eggs laid per worm, indicating that the assay was functional (S2 Fig; S1 Table, row 6). Statistical analysis revealed no effect of DOI dosage on the number of eggs laid per worm (Fig 3; S1 Table, row 7). We conclude that DOI has no detectable effect on egg-laying under our conditions. We note that LSD has no effect on egg-laying in *Drosophila* [19].

**Fig 3.**
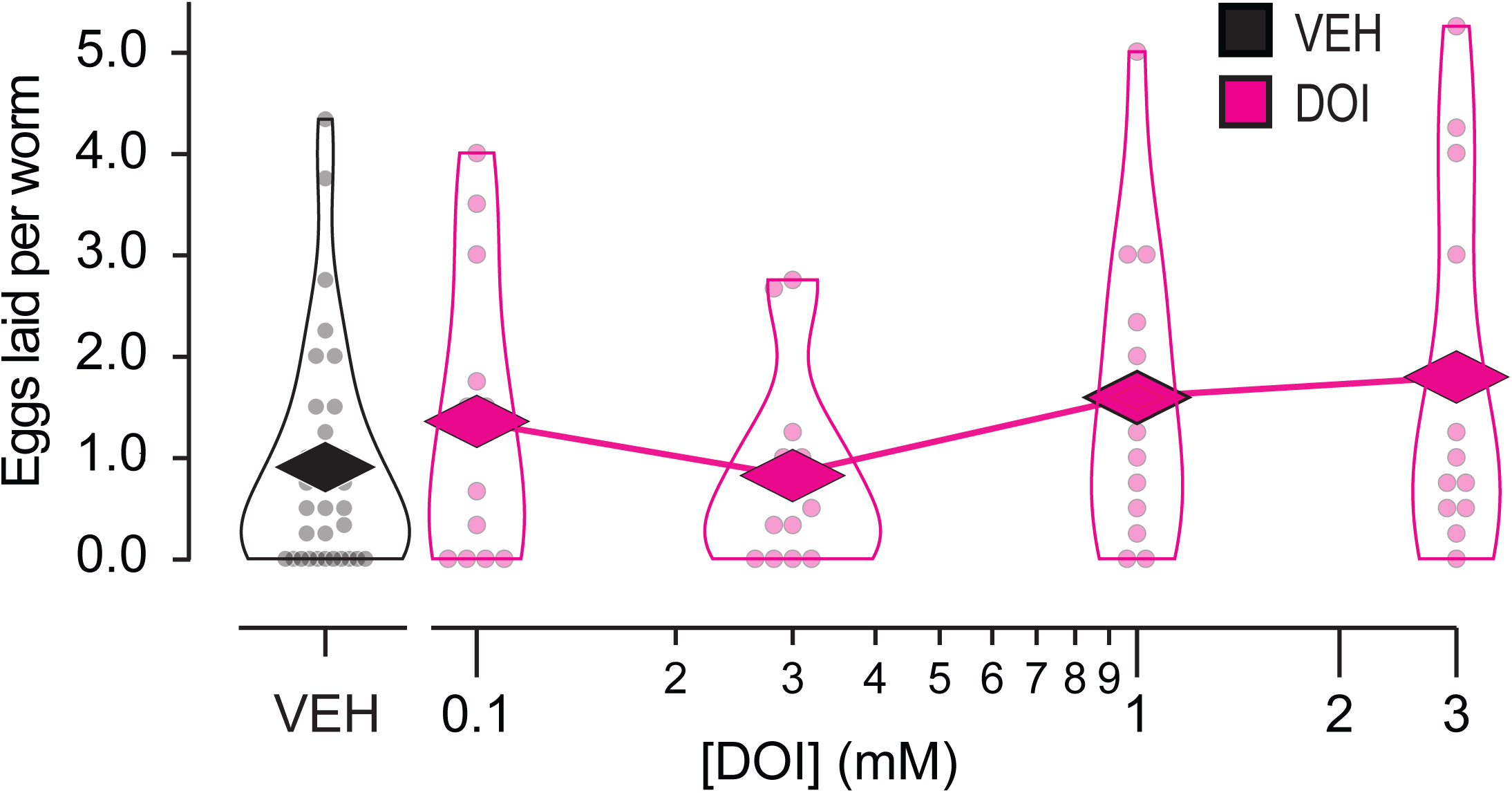
Absence of Effect of DOI on egg laying. Mean number of eggs laid per worm is plotted against DOI concentration. Worms were exposed to DOI for 30 min while they laid eggs. Each data point represents a cohort of 10-15 worms. *n* = 12 cohorts for DOI at each concentration and 48 for VEH (vehicle). *Diamonds*, means. See S1 Table, row 7.

### DOI Suppresses Feeding

Worms ingest bacterial food through rhythmic contractions of the pharynx, a tube-shaped muscular organ that delivers bacterial particles to the gut. Each pharyngeal contraction is called a pump. To test whether DOI has an effect on pumping, we pre-exposed young adult worms to M9 solutions containing food, to simulate pumping, and DOI at various concentrations (in mM: 0.1, 0.3, 1.0, 3.0) for 30 minutes; control worms were exposed to the food solution only. After pre-exposure, worms, along with their exposure solution, were injected into a microfluidic device fitted with electrodes that recorded the electrical signals associated with each pharyngeal contraction [28]. Such recordings are called electropharyngeograms (EPGs). EPGs from individual worms were recorded for 5 minutes each; these recording were made in close succession over a 90-minute period that began when worms were injected into the recording device. At doses exceeding 0.1 mM, DOI suppressed pharyngeal pumping in a dose-dependent manner (Fig 4A; S1 Table, row 8). This result is reminiscent of DOI-induced hypophagia in rodents [37].

**Fig 4.**
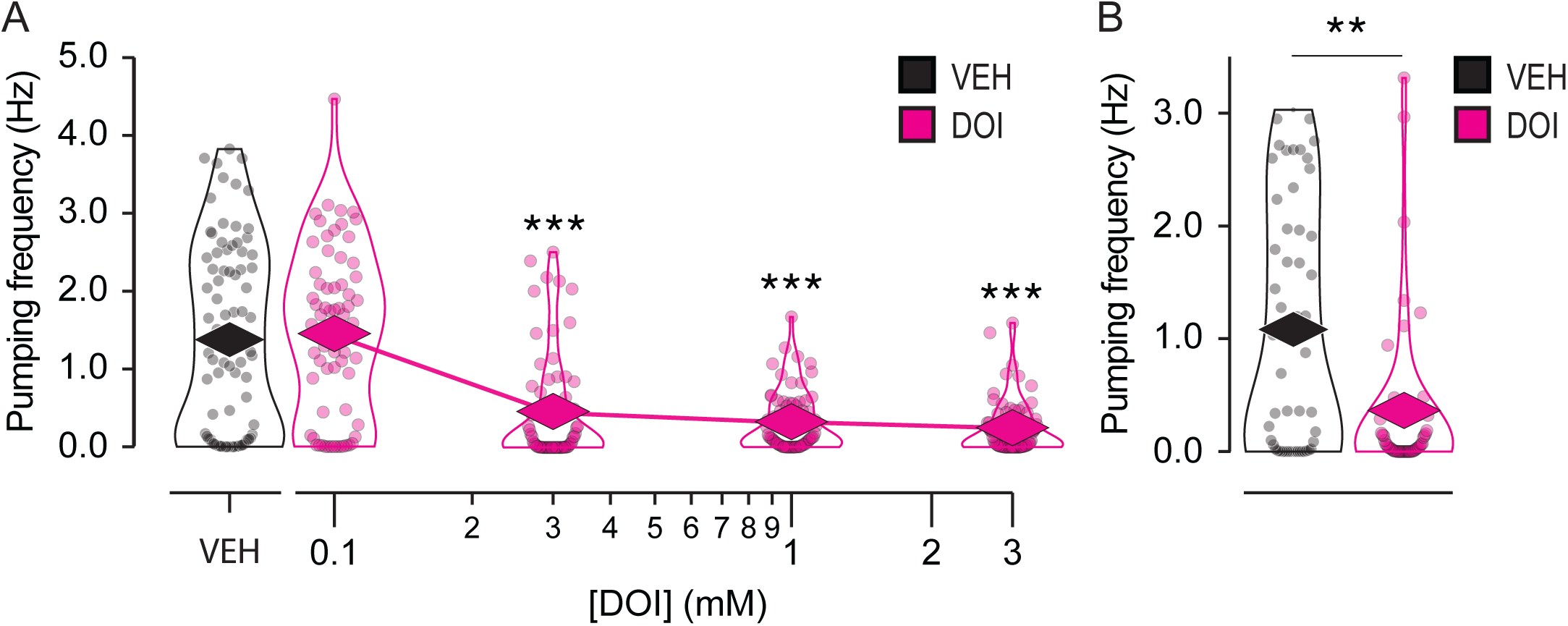
Effect of DOI on feeding rate. . A. Pharyngeal pumping rate is plotted against DOI concentration. Worms were exposed to DOI for 30–120 minutes, depending on the order in which they were recorded. Each data point represents a single worm. *n* = 65 for DOI and 84 for VEH (vehicle). ***, Wilcoxon rank sum test, significantly different from VEH; *p* < 6.33 × 10^-7^. See S1 Table, row 8. B. Effect of 1 mM DOI on feeding rate in a mutant that lacks all chemosensory neuron function (Table 1, XL344). Each data point represents a single worm. *n* = 55 worms for DOI and VEH. **, Wilcoxon rank sum test, *p* = 1.80 × 10^-3^. See S1 Table, row 11. *Diamonds*, means.

It is notable that DOI has a strong effect on pumping but no discernable effects on crawling, swimming, or egg laying under our conditions. One difference between the pumping assay and the others is that the worms were recorded over a period of 90 minutes after pre-exposure (see Materials and Methods), thus all worms were exposed to DOI for a longer period than in any of the other assays. Perhaps only those worms that were recorded later in the assay showed suppression. To test whether this was the case, we plotted mean pumping frequency against the order in which each worm was recorded, thereby using order as a proxy for time (S3 Fig). This was done by plotting mean pumping rate as a function of recording order at a DOI concentration of 1 mM (S3 Fig). Statistical analysis showed no effect of recording order on pumping rate and no interaction between recording order and effect of DOI (S1 Table, row 9). Thus, suppression was stable throughout the recording period. Furthermore, the effect of DOI was detectable in the first worm recorded in each session (S1 Table, row 10). This analysis shows that pumping suppression was present immediately after the pre-exposure period and constant thereafter.

We conclude that DOI’s strong effect on pumping is not attributable to the longer overall exposure period in this assay (≤120 minutes).

It is possible that DOI suppresses pumping because it has an aversive taste or smell. For example, the bitter substance quinine suppresses pumping [38]. Accordingly, we tested the effect of DOI on a mutant strain in which the function of all taste and olfactory neurons was eliminated by a mutation that prevents the formation of chemosensory cilia [39]. At a DOI concentration of 1 mM, pumping was strongly suppressed in the mutant relative to vehicle-only controls (Fig 4B; S1 Table, row 11). This result indicates that pumping inhibition is probably not a response to possible aversive chemosensory properties of DOI solutions.

### Feeding Suppression Is Intact in Serotonin Receptor Mutants

In mammals, DOI has high affinity for serotonin receptors. To date, DOI is known to bind to two serotonin receptors in *C. elegans*, SER-1 and SER-4, with binding constants similar to that of serotonin [40,41]. Although serotonin receptors are associated with the stimulation of pumping, they could, in principle, suppress pumping by excitation block which has been reported in the pharynx [42]. Therefore, we asked whether any of the six *C. elegans* serotonin receptor genes were necessary for DOI-induced suppression of pharyngeal pumping.

Knockout mutants for each serotonin receptor (SER-1, SER-4, SER-5, SER-7, MOD-1, LGC-50) were exposed to DOI and recorded as described above, now at a single DOI concentration (1 mM). There were significant main effects of strain and drug, the latter implying that suppression persisted in the mutants overall (Fig 5; S1 Table, row 12). Post hoc tests comparing each mutant to the N2 reference strain revealed main effects of drug but no drug × strain interactions, suggesting that the suppressive effect of DOI is similar in the mutants and reference strain (S1 Table, rows 13-18). We conclude that serotonin receptor genes are not individually required for DOI-induced pumping suppression. This result rules out a model in which activation of a single type of serotonin receptor by DOI is necessary and sufficient for pumping suppression. However, it does not exclude a model in which multiple serotonin receptors act redundantly to suppress pumping. In related work, the receptors SER-1 and SER-4 have been shown to be required for LSD-induced locomotory slowing [22].

**Fig 5.**
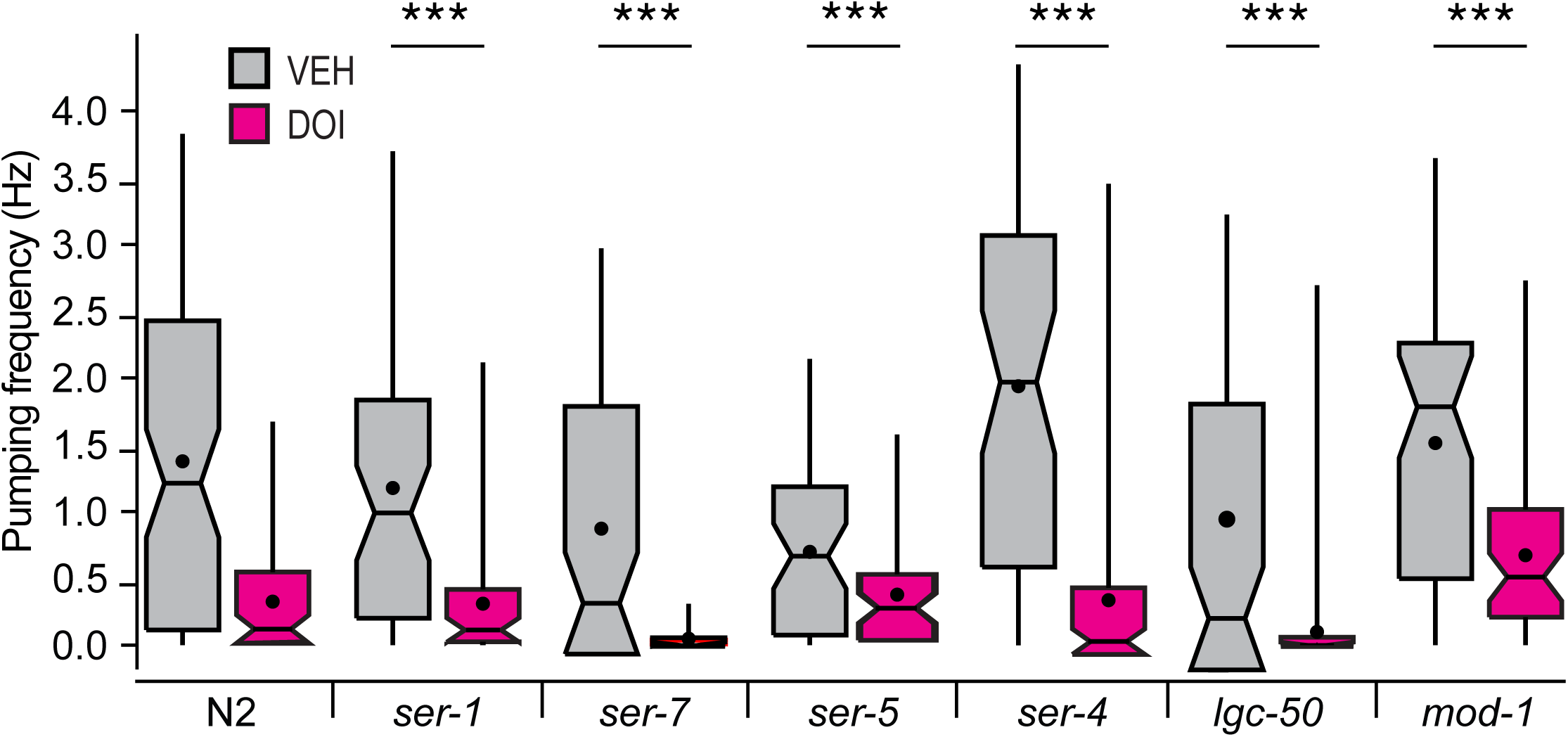
Effect of DOI on feeding rate in serotonin receptor mutants. . Pumping frequency ± DOI is shown for the reference strain N2 and knockout mutant strains for each of the worm’s six serotonin receptor genes. Worms were exposed to 1 mM DOI for 30-120 minutes, depending on the order in which they were recorded. *n* ≥ 55 worms for DOI and VEH (vehicle). Brunner-Munzel Test revealed significant main effects of strain and DOI and a significant strain × DOI interaction. See S1 Table, row 12. ***, Post hoc Brunner-Munzel test, N2 versus each mutant, main effect of DOI, *p* < 3.72 × 10^-8^ (S1 Table, rows 13-18). *Box*, 1^st^ and 3^rd^ quartiles; *notch*, median; *dot*, mean; *whiskers*, data range.

## Discussion

In this study, one of the first to investigate the effects of psychedelics on *C. elegans* behavior [22], we performed a preliminary characterization of the effects of DOI on four characteristic behaviors. Under our conditions, there was no detectable effect of DOI on crawling, swimming, and egg laying, but a profound suppressive effect on pharyngeal pumping. Thus, DOI has a behavioral phenotype that can now be utilized in mode of action studies.

### Limitations

The main limitation of this study is the difficulty of interpreting negative results in *C. elegans* pharmacology. While positive results—such as those from the toxicity (Fig 1) and pharyngeal pumping assays (Figs 4, 5)—confirm DOI absorption, negative results (Figs 2, 3) could reflect insufficient drug uptake. In a liquid environment, *C. elegans* can absorb drugs passively via diffusion through the cuticle and actively via pharyngeal pumping [43], but immersion alone does not trigger pumping unless food or serotonin is present [44]. Several findings support passive absorption of DOI in *C. elegans*: (i) It is a common delivery route for small molecules like DOI (358 g/mol) [45–47]; (ii) In the toxicity assay, DOI was effective without a pumping stimulant, leaving diffusion as the sole delivery route; (iii) LSD (323 g/mol) is passively absorbed by *C. elegans* as shown by high-pressure liquid chromatography–mass spectrometry [22]. Still, DOI may require higher doses, longer exposures, or more sensitive assays for its effects on locomotion and egg-laying to be revealed, such as high-resolution locomotion tracking [48,49] or testing in hypotonic media which enhance serotonergic responses [50,51]. Nevertheless, whereas we cannot rule out DOI effects on locomotion and egg laying entirely, our data show no detectable impact on these behaviors under standard conditions.

### Toxicity

We found DOI to be benign at low to moderate doses and toxic at the highest dose. Exposing worms to 10 mM DOI for 30 minutes reduced 24-hour survival rate from 95% to 20%. There appear to be no reports of DOI toxicity in other organisms. However, DOI (10-100 μM) produces cell death in cultured rat embryonic cortical and hippocampal neurons [52,53]. The PubChem database [54] does not list the LD50 for DOI. However, the LD50 for 2,5-dimethoxy-4-methylamphetamine (DOM), which is structurally almost identical to DOI, is 94 mg/kg (mouse; intraperitoneal) [55] or about 310 μM (see Materials and Methods for assumptions and conversion equation). As for *C. elegans*, the worm’s cuticle is known to be a significant diffusion barrier so the actual internal concentration of externally applied drugs is probably much lower [33].

Although the ability of *C. elegans* to absorb DOI is unknown, its ability to absorb LSD has been measured as function of external LSD concentration for a 15 minute exposure [22]. These measurements suggest that the internal concentration is 30–200-fold lower than the external concentration. Thus, If DOI is absorbed like LSD, exposure to 10 mM DOI would result in an internal concentration of 50–300 μM. The upper limit of this range is similar to the expected internal concentration in mice given the LD50 dose of DOM (see above). These findings suggest that DOI may exhibit organismal toxicity at concentrations comparable to the known LD50 of structurally related compounds.

### Biological relevance

A related question is whether the non-toxic DOI concentrations we used (0.1–3 mM) are in a biologically relevant concentration range. In studies using rodent models, DOI dosage is on the order of 0.1-10 mg/kg [56]. Converted to molar units (see Materials and Methods for assumptions and conversion equation), the dosage range is 0.33-33 μM. In *C. elegans*, LSD is passively absorbed to an internal concentration that is is 30–200-fold less than the external concentration [22]. Assuming DOI diffuses like LSD, a mid-range external concentration of 1 mM DOI would result in an internal concentration of 5–30 μM, which overlaps with the range used in rodent studies. This reasoning suggests that the non-toxic DOI concentrations we used are probably biologically relevant.

### Pumping suppression

The inhibitory effect of DOI on pumping is surprising. As DOI is a potent serotonin receptor agonist, and serotonin stimulates pumping [57], one might have expected DOI to stimulate pumping. Previous findings support a simple model to explain DOI’s inhibitory effect on pumping.

Pumping suppression is part of the normal regulation of pharyngeal activity in which excitatory and inhibitory pathways converge on the pharynx and are likely active simultaneously to ensure that worms pump only under appropriate circumstances [38,58]. Excitatory pathways facilitate pumping in response to bacteria or odors that predict them [38,59], whereas inhibitory pathways suppress pumping in response to the absence of bacteria or the presence of aversive compounds such as quinine [38,58]. We propose a model in which DOI acts on one or more receptors involved in the worm’s native pathways for pumping suppression. Components of several inhibitory pathways have been identified. The biogenic amines octopamine and tyramine suppress serotonin-stimulated pumping when applied exogenously [57,60–62]. Thus, receptors for tyramine–SER-2, TYRA-2, TYRA-3, LGC-55 [63–66]– and octopamine–SER-3, SER-6, and OCTR-1 [65,67,68]–are potential DOI targets. Tyramine receptors in *C. elegans*, though sensitive to tyramine, are orthologous to serotonin receptors in mammals [69], whereas octopamine receptors are orthologous to adrenergic receptors, to which DOI also binds [70]. The model can be tested by investigating the effect of DOI on pumping in tyramine and octopamine receptor mutants.

If tyramine or octopamine receptors are required for pumping suppression, the model must account for the fact that tyramine and octopamine also modulate locomotion and egg laying yet modulation of these behaviors was not observed in our study. Exogenous tyramine immobilizes crawling worms [63], whereas octopamine activates a “roaming” state, characterized by extended bouts of relatively fast forward movement; roaming is a response to food depletion [71–74]. Both amines inhibit egg laying [57,75–77]. One simple possibility is that pharyngeal pumping is more sensitive to DOI than locomotion or egg laying. Alternatively, it is possible that we failed to detect DOI effects on these behaviors because worms were in a state that was not conducive to this type of modulation. For example, because worms were exposed in the absence of food, and were tested while crawling on a foodless substrate, they may have been in the roaming state much of the time, thereby occluding an increase in speed. Similarly, in the egg laying assay worms may have been in a low egg-laying state because they were tested in the absence of food or serotonin [57]. Further research is needed to test this model.

An alternative model involves modulation of pathways that stimulate pumping. Another class of psychoactive drugs, antipsychotics, can suppress pumping in *C. elegans* [42,78]. Examples include trifluoperazine, fluphenazine, clozapine, and olanzapine. In the case of clozapine, pumping suppression is reduced in knockouts of the nicotinic acetylcholine receptor ACR-7, which is expressed on pharyngeal muscles. Furthermore, the effects of *acr-7* knockouts can be mimicked by acetylcholine receptor antagonists [42]. Therefore, clozapine is believed to silence the pharynx through direct activation of ACR-7, causing tetanic contraction of pharyngeal muscles similar to that observed upon bath application of nicotine [79]. Thus, a possible mode of action of DOI could be overstimulation of excitatory pathways.

### Related work

Several other studies had assessed the effects of psychedelics on genetically tractable organisms. In *C. elegans*, LSD reduces crawling speed [22], an effect that requires *ser-1* and *ser-4*. Work on *Drosophila* is more advanced. LSD appears not to modulate egg laying [19], but it reduces locomotor activity and impairs performance on an optomotor task [20]. Psilocybin causes decreased immobility in the Drosophila equivalent of the forced swim test, a model for behavioral despair. DOI decreases the overall level of aggressive behavior in a manner that requires *Drosophila* orthologs of 5-HT2 receptors [80]. These studies, together with the present one, underscore the utility of genetically tractable organisms in the behavioral and genetic analysis of psychedelic action.

## Acknowledgements

Supported by National Institute of General Medical science (R35 GM152169). Some strains were provided by the CGC, which is funded by NIH Office of Research Infrastructure Programs (P40 OD010440).

## Supporting information captions

**S1 Fig. Crawling speed versus time at the indicated dose of DOI.** Each data point represents the mean speed in 30 sec. bins across cohorts of 10-15 worms. *n* = 8 cohorts at each dose of DOI and 31 for VEH (vehicle). DOI exposure time was 30 min. Error bars ± 95% CI. *Shaded gray trace*, VEH. Two-way ANOVA, dose versus time, main effect of time, *p* =9.86 × 10^-34^, no effect of DOI dose, and no dose × time interaction. See S1 Table, row 4.

**S2 Fig. Positive control for the egg-laying assay using serotonin (5-HT) as a stimulant.**The assay was carried out on cohorts of 10–15 worms. *n* = 8 cohorts for DOI and VEH (vehicle). Cohorts were exposed to 5-HT for 30 min. at 5 mg/mL. *Diamonds*, means; ***, two-sample *t*-test, *p* = 5.16 x 10^-4^. See S1 Table, row 6.

**S3 Fig. Mean pumping frequency versus the order in which each worm was recorded.**Same experiment as in Fig 4A. Recording order serves as a proxy for time in the recording device.

Worms were exposed to DOI for 30-120 minutes, depending on the order in which they were recorded. Each point represents the mean pumping frequency for individual worms recorded at a particular rank in a series of 12 recordings. Error bars, ± CI; **, two-sample *t*-test, *p* = 0.016; •••, Two-way ANOVA order versus DOI, main effect of DOI, *p* = 6.16 × -10^-10^, no effect of order, no order × DOI interaction. See S1 Table, row 9,10.

**S1 Table. Details of statistical analyses.** *n* indicates that the unit of replication was single worms; *n*\* indicates the unit of replication was a cohort of 10-15 worms. The notation e^-x^ represents 10^-x^. Significant *p* values (*p* < 0.05) are in bold text. *Symbols*: **, *p* < 0.01; ***, *p* < 0.001; †, *p* < 0.05 after Bonferroni correction; †††, *p* < 0.001 after Bonferroni correction.

**S2 Table. Power analysis of crawling, swimming, and egg-laying assays.** The terms *N1* and *N2* are the sample size in the experiment. Power is the likelihood of detecting an effect if the effect exists. The *minimum detectable effect size* (d_min_) represents the smallest true effect that the experimental design is likely to detect with the specified power, significance level, and sample size. We used a significance level (α) of 0.05 and a power (1 – β) of 0.8, where β is the probability of a false negative result. Sample sizes are given in the table. The minimum detectable effect size is expressed in a Cohen’s *d* framework,

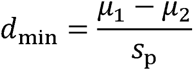

where µ_1_and µ_2_ are the means of two samples (control and drug) and s_p_ is the *empirical pooled standard deviation* observed in the actual experiment. The quantity d_min_ was computed using G*Power [30]. The minimum detectable difference between means was computed as (µ_1_ - µ_2_) = d_min_ · s_p_. Minimum detectable effect sizes and differences between means were computed using s_p_ from: vehicle versus 0.1 mM DOI (Fig 2A), vehicle versus DOI at *t* = 30 min. (Fig 2B), and vehicle versus 0.1 mM DOI (Fig 3) as typical examples.

**S1 File. All data acquired in this study.**

## Notes

### Competing Interest Statement

S.R.L. declares a competing financial interest in InVivo Biosystems, Inc., which manufactures the ScreenChip system.

### Summary of Updates

In response to concerns of reviewers, we added the effect of DOI on swimming and an analysis of the time course of its effect on crawling. Also, the Discussion was completely rewritten to modify claims made in the paper.

