## Supplemental Table 1 for "Behavioral and genetic analysis of the effects of the psychedelic 2,5-dimethoxy-4-iodoamphetamine (DOI) in *C. elegans*"

| **Fig.** | **Row** | **Comparison** | **Test** | **Number of replicates** | | **Mean ± 95% CI** | **Statistic** | **p-value** | **Signif.** | **Method** | **Effect size** | |
| --- | --- | --- | --- | --- | --- | --- | --- | --- | --- | --- | --- | --- |
| 1 | 1 | DOI effect on survival in N2: VEH vs [DOI] | ANOVA | VEH, *n** = 15 | | 92.89 ± 4.94 |  |  |  |  |  | |
|  |  |  |  | DOI 0.1 mM, *n** = 3 | | 94.32 ± 14.64 |  |  |  |  |  |  |
|  |  |  |  | DOI 0.3 mM, *n** = 3 | | 95.16 ± 11.83 |  |  |  |  |  |  |
|  |  |  |  | DOI 1 mM, *n** = 3 | | 90.24 ± 6.14 |  |  |  |  |  |  |
|  |  |  |  | DOI 3 mM, *n** = 3 | | 93.63 ± 14.24 |  |  |  |  |  |  |
|  |  |  |  | DOI 10 mM, *n** = 3 | | 12.74 ± 27.74 |  |  |  |  |  |  |
|  |  |  | Main effect of DOI Dose |  | |  | *F*(5,24) = 53.88 | **2.9e^-12^** | *** |  |  | |
|  | **Post-hoc tests** | | | | | | | | | | | |
|  | 2 | [DOI] 0 vs 0.1 mM | Dunnett's test |  | |  | *q* = -0.28 | 0.971 |  |  |  | |
|  |  | [DOI] 0 vs 0.3 mM |  |  | |  | *q* = -0.45 | 0.983 |  |  |  | |
|  |  | [DOI] 0 vs 1 mM |  |  | |  | *q* = 0.52 | 0.768 |  |  |  | |
|  |  | [DOI] 0 vs 3 mM |  |  | |  | *q* = -0.15 | 0.955 |  |  |  | |
|  |  | [DOI] 0 vs 10 mM |  |  | |  | *q* = 15.74 | **8.0e^-7^** | *** | Cohen's d | -7.93 | |
| 2A | 3 | DOI effect on crawling speed in N2, VEH vs [DOI] | ANOVA | VEH, *n** = 31 | | 136.80 ± 12.85 |  |  |  |  |  | |
|  |  |  |  | DOI 0.1 mM, *n** = 8 | | 135.12 ± 26.38 |  |  |  |  |  |  |
|  |  |  |  | DOI 0.3 mM, *n** = 8 | | 151.05 ± 23.43 |  |  |  |  |  |  |
|  |  |  |  | DOI 1 mM, *n** = 8 | | 140.99 ± 22.09 |  |  |  |  |  |  |
|  |  |  |  | DOI 3 mM, *n** = 8 | | 127.71 ± 23.40 |  |  |  |  |  |  |
|  |  |  | Main effect of DOI Dose |  | |  | *F*(4,58) = 0.58 | 0.678 |  |  |  | |
| S1 | 4 | DOI effect on crawling time course: [DOI] vs VEH | 2-way mixed ANOVA | | VEH, *n** = 31 |  |  |  |  |  |  | |
|  |  |  |  |  | DOI 0.1 mM, *n** = 8 |  |  |  |  |  |  |  |
|  |  |  |  |  | DOI 0.3 mM, *n** = 8 |  |  |  |  |  |  |  |
|  |  |  |  |  | DOI 1 mM, *n** = 8 |  |  |  |  |  |  |  |
|  |  |  |  |  | DOI 3 mM, *n** = 8 |  |  |  |  |  |  |  |
|  |  |  | Main effect of Time | |  |  | *F*(4.70, 272.7)=53.458 | **9.86e^-34^** | *** | Eta-squared | 0.306 | |
|  |  |  | Main effect of [DOI] | |  |  | *F*(4,58)= 1.495 | 0.346 |  |  |  | |
|  |  |  | Interaction, Time × [DOI] | |  |  | *F*(18.81,272.7)=0.905 | 0.536 |  |  |  | |
| **Fig.** | **Row** | **Comparison** | **Test** | **Number of replicates** | | **Mean ± 95% CI** | **Statistic** | **p-value** | **Signif.** | **Method** | **Effect size** | |
| 2B | 5 | DOI effect on swimming wave initiation frequency: DOI vs VEH | 2-way mixed ANOVA | Baseline  VEH, *n* = 90  DOI 1 mM, *n* = 97 | | 92.71 ± 6.57  97.40 ± 4.44 |  |  |  |  | |  |
|  |  |  |  | t=0  VEH, *n* = 84  DOI 1 mM, *n* = 97 | | 89.76 ± 5.22  92.52 ± 3.36 |  |  |  |  |  |  |
|  |  |  |  | T=30  VEH, *n* = 88  DOI 1 mM, *n* = 98 | | 91.46 ± 5.43  89.13 ± 3.81 |  |  |  |  |  |  |
|  |  |  | Main effect of Time |  | |  | *F*(1.93,355.75) =5.560 | **0.005** | ** | eta-squared | | 0.01 |
|  |  |  | Main effect of DOI |  | |  | *F*(1,184) = 0.065 | 0.799 |  |  | |  |
|  |  |  | Interaction, Time × DOI |  | |  | *F*(1.93,355.75)=0.812 | 0.441 |  |  | |  |
| S2 | 6 | Effect of 5-HT on egg-laying: VEH vs 5-HT | *t*-test | VEH, *n** = 8, 5-HT, *n** = 8 | | VEH: 0.53 ± 0.70 5-HT: 3.39 ± 1.21 | *t*(11.20) =-4.81 | **0.001** | *** | Cohen's d | 2.4 | |
| 3 | 7 | DOI effect on egg laying in N2: VEH vs [DOI] | ANOVA | VEH, *n** = 8 | | 0.90 ± 0.40 |  |  |  |  |  | |
|  |  |  |  | DOI 0.1 mM, *n** = 8 | | 1.35 ± 0.93 |  |  |  |  |  |  |
|  |  |  |  | DOI 0.3 mM, *n** = 8 | | 0.82 ± 0.63 |  |  |  |  |  |  |
|  |  |  |  | DOI 1 mM, *n** = 8 | | 1.59 ± 0.97 |  |  |  |  |  |  |
|  |  |  |  | DOI 3 mM, *n** = 8 | | 1.79 ± 1.16 |  |  |  |  |  |  |
|  |  |  | Main effect of DOI Dose |  | |  | *F*(4,75) = 1.51 | 0.208 |  |  |  | |
| 4A | 8 | DOI effect on pumping frequency in N2: VEH vs [DOI] | Wilcoxon rank sum test | VEH, *n* = 84, DOI 0.1 mM, *n* = 65 | | VEH: 1.38 ± 0.26 DOI: 1.45 ± 0.27 | *z* = -0.39 | 0.653 |  |  |  | |
|  |  |  |  | VEH, *n* = 84, DOI 0.3 mM, *n* = 65 | | VEH: 1.38 ± 0.26 DOI: 0.46 ± 0.17 | *z* = 4.99 | **2.84e^-7^** | ***/††† | rank-serial | -0.48 | |
|  |  |  |  | VEH, *n* = 84, DOI 1 mM, *n* = 65 | | VEH: 1.38 ± 0.26 DOI: 0.33 ± 0.10 | *z* = 4.84 | **6.33e^-7^** | ***/††† | rank-serial | -0.46 | |
|  |  |  |  | VEH, *n* = 84, DOI 3 mM, *n* = 65 | | VEH: 1.38 ± 0.26 DOI: 0.25 ± 0.09 | *z* = 5.56 | **1.24e^-8^** | ***/††† | rank-serial | -0.53 | |
| **Fig.** | **Row** | **Comparison** | **Test** | **Number of replicates** | | **Mean ± 95% CI** | **Statistic** | ***p-*value** | **Signif.** | **Effect size measure** | **Effect size** | |
| S3 | 9 | Effect of DOI and recording order on EPG frequency | 2-way ANOVA | VEH, *n* = 84  DOI 1 mM, *n* = 65 | |  |  |  |  |  | |  |
|  |  |  | Main effect of Order |  | |  | *F*(1,145)= 0.904 | 0.343 |  |  | |  |
|  |  |  | Main effect of DOI |  | |  | *F*(1,145)= 43.964 | **6.16e^-10^** | *** | eta-squared | | 0.23 |
|  |  |  | Interaction, Order × DOI |  | |  | *F*(1,145)= 0.018 | 0.895 |  |  | |  |
|  | **Post-hoc tests** | | | | | | | | | | | |
|  | 10 | First recording in a session:  VEH vs DOI 1 mM | Welch Two Sample *t*-test | VEH, *n* = 9  DOI 1 mM, *n* = 5 | | 1.187 ± 0.24   0.20 ± 1.73 | *t*(9.53)= 2.980 | **0.0145** | ** | Cohen’s d | | 1.26 |
| 4B | 11 | DOI effect on pumping frequency in a chemosensory-blind strain (XL344): VEH vs DOI | Wilcoxon rank sum test | XL344, VEH, *n* = 55 | | 1.08 ± 0.29 | *z* = 3.12 | **1.8e^-3^** | ** | rank-serial | -0.35 | |
|  |  |  |  | XL344, DOI 1 mM, *n* = 55 | | 0.32 ± 0.18 |  |  |  |  |  |  |
| 5 | 12 | DOI effect on pumping frequency in serotonin receptor mutants: N2 vs mutants | Brunner-Munzel Test ANOVA-type statistics | N2, VEH, *n* = 84 | | 1.38 ± 0.26 |  |  |  |  |  | |
|  |  |  |  | N2, DOI 1 mM, *n* = 65 | | 0.33 ± 0.10 |  |  |  |  |  |  |
|  |  |  |  | *ser1(ok345)*, VEH, *n* = 55 | | 1.18 ± 0.29 |  |  |  |  |  |  |
|  |  |  |  | *ser1(ok345)*, DOI 1 mM, *n* = 55 | | 0.31 ± 0.12 |  |  |  |  |  |  |
|  |  |  |  | *ser-7(tm1325)*, VEH, *n* = 55 | | 0.87 ± 0.25 |  |  |  |  |  |  |
|  |  |  |  | *ser-7(tm1325)*, DOI 1 mM, *n* = 55 | | 0.05 ± 0.02 |  |  |  |  |  |  |
|  |  |  |  | *ser-5(vq1)*, VEH, *n* = 55 | | 1.94 ± 0.38 |  |  |  |  |  |  |
|  |  |  |  | *ser-5(vq1)*, DOI 1 mM, *n* = 55 | | 0.34 ± 0.17 |  |  |  |  |  |  |
|  |  |  |  | *ser-4(ok512)*, VEH, *n* = 55 | | 0.70 ± 0.17 |  |  |  |  |  |  |
|  |  |  |  | *ser-4(ok512)*, DOI 1 mM, *n* = 55 | | 0.38 ± 0.11 |  |  |  |  |  |  |
|  |  |  |  | *lgc-50(tm3712)*, VEH, *n* = 55 | | 0.94 ± 0.29 |  |  |  |  |  |  |
|  |  |  |  | *lgc-50(tm3712)*, DOI 1 mM, *n* = 55 | | 0.10 ± 0.10 |  |  |  |  |  |  |
|  |  |  |  | *mod-1(ok103)*, VEH, *n* = 55 | | 1.51 ± 0.29 |  |  |  |  |  |  |
|  |  |  |  | *mod-1(ok103)*, DOI 1 mM, *n* = 55 | | 0.67 ± 0.16 |  |  |  |  |  |  |
|  |  |  | Main effect of Strain |  | |  | *F*(5.86,665.25)= 15.63 | **9.23e^-17^** | *** |  |  | |
|  |  |  | Main effect of DOI |  | |  | *F*(1,665.25) = 181.04 | **1.12e^-36^** | *** |  |  | |
|  |  |  | Interaction, Strain × DOI |  | |  | *F*(5.86,665.25) = 3.89 | **9e^-4^** | *** |  |  | |
|  | **Post-hoc tests** | | | | | | | | | | | |
|  | 13 | DOI effect on pumping frequency: N2 vs ser1(ok345) ± DOI 1 mM | Brunner-Munzel Test ANOVA-type statistics | N2, VEH, *n* = 84 | | 1.38 ± 0.26 |  |  |  |  |  | |
|  |  |  |  | N2, DOI 1 mM, *n* = 65 | | 0.33 ± 0.10 |  |  |  |  |  |  |
|  |  |  |  | *ser1(ok345)*, VEH, *n* = 55 | | 1.18 ± 0.29 |  |  |  |  |  |  |
|  |  |  |  | *ser1(ok345)*, DOI 1 mM, *n* = 55 | | 0.31 ± 0.12 |  |  |  |  |  |  |
|  |  |  | Main effect of Strain |  | |  | *F*(1,234.96) = 0.34 | 0.561 |  |  |  | |
|  |  |  | Main effect of DOI |  | |  | *F*(1,234.96) = 57.84 | **6.80e^-13^** | ***/††† | Cliff’s Δ | -0.49 | |
|  |  |  | Interaction, Strain × DOI |  | |  | *F*(1,234.96) = 0.12 | 0.728 |  |  |  | |
|  | 14 | DOI effect on pumping frequency: N2 vs ser-7(tm1325) ± DOI 1 mM | Brunner-Munzel Test ANOVA-type statistics | N2, VEH, *n* = 84 | | 1.38 ± 0.26 |  |  |  |  |  | |
|  |  |  |  | N2, DOI 1 mM, *n* = 65 | | 0.33 ± 0.10 |  |  |  |  |  |  |
|  |  |  |  | *ser-7(tm1325)*, VEH, *n* = 55 | | 0.87 ± 0.25 |  |  |  |  |  |  |
|  |  |  |  | *ser-7(tm1325)*, DOI 1 mM, *n* = 55 | | 0.05 ± 0.02 |  |  |  |  |  |  |
|  |  |  | Main effect of Strain |  | |  | *F*(1,200.77) = 26.48 | **6.32e^-07^** | ***/††† | Cliff’s Δ | -0.34 | |
|  |  |  | Main effect of DOI |  | |  | *F*(1,200.77) = 55.24 | **3.01e^-12^** | ***/††† | Cliff’s Δ | -0.48 | |
|  |  |  | Interaction, Strain × DOI |  | |  | *F*(1,200.77) = 1.34 | 0.249 |  |  |  | |
|  | 15 | DOI effect on pumping frequency: N2 vs ser-5(vq1) ± DOI 1 mM | Brunner-Munzel Test ANOVA-type statistics | N2, VEH, *n* = 84 | | 1.38 ± 0.26 |  |  |  |  |  | |
|  |  |  |  | N2, DOI 1 mM, *n* = 65 | | 0.33 ± 0.10 |  |  |  |  |  |  |
|  |  |  |  | *ser-5(vq1)*, VEH, *n* = 55 | | 1.94 ± 0.38 |  |  |  |  |  |  |
|  |  |  |  | *ser-5(vq1)*, DOI 1 mM, *n* = 55 | | 0.34 ± 0.17 |  |  |  |  |  |  |
|  |  |  | Main effect of Strain |  | |  | *F*(1,208.87) = 0.15 | 0.702 |  |  |  | |
|  |  |  | Main effect of DOI |  | |  | *F*(1,208.87) = 82.36 | **8.60e^-17^** | ***/††† | Cliff’s Δ | -0.55 | |
|  |  |  | Interaction, Strain × DOI |  | |  | *F*(1,208.87) = 5.98 | 0.015 | ** |  |  | |
|  | 16 | DOI effect on pumping frequency: N2 vs ser-4(ok512) ± DOI 1 mM | Brunner-Munzel Test ANOVA-type statistics | N2, VEH, *n* = 84 | | 1.38 ± 0.26 |  |  |  |  |  | |
|  |  |  |  | N2, DOI 1 mM, *n* = 65 | | 0.33 ± 0.10 |  |  |  |  |  |  |
|  |  |  |  | *ser-4(ok512)*, VEH, *n* = 55 | | 0.70 ± 0.17 |  |  |  |  |  |  |
|  |  |  |  | *ser-4(ok512)*, DOI 1 mM, *n* = 55 | | 0.38 ± 0.11 |  |  |  |  |  |  |
|  |  |  | Main effect of Strain |  | |  | *F*(1,239.45) = 1.35 | 0.246 |  |  |  | |
|  |  |  | Main effect of DOI |  | |  | *F*(1,239.45) = 32.36 | **3.72e^-08^** | ***/††† | Cliff’s Δ | -0.4 | |
|  |  |  | Interaction, Strain × DOI |  | |  | *F*(1,239.45) = 3.64 | 0.058 |  |  |  | |
|  | 17 | DOI effect on pumping frequency: N2 vs lgc-50(tm3712) ± DOI 1 mM | Brunner-Munzel Test ANOVA-type statistics | N2, VEH, *n* = 84 | | 1.38 ± 0.26 |  |  |  |  |  | |
|  |  |  |  | N2, DOI 1 mM, *n* = 65 | | 0.33 ± 0.10 |  |  |  |  |  |  |
|  |  |  |  | *lgc-50(tm3712)*, VEH, *n* = 55 | | 0.94 ± 0.29 |  |  |  |  |  |  |
|  |  |  |  | *lgc-50(tm3712)*, DOI 1 mM, *n* = 5555 | | 0.10 ± 0.10 |  |  |  |  |  |  |
|  |  |  | Main effect of Strain |  | |  | *F*(1,194.10) = 24.34 | **1.73e^-06^** | ***/††† | Cliff’s Δ | -0.34 | |
|  |  |  | Main effect of DOI |  | |  | *F*(1,194.10) = 47.62 | **7.14e^-11^** | ***/††† | Cliff’s Δ | -0.46 | |
|  |  |  | Interaction, Strain × DOI |  | |  | *F*(1,194.10) = 0.94 | 0.334 |  |  |  | |
|  | 18 | DOI effect on pumping frequency: N2 vs mod-1(ok103) ± DOI 1 mM | Brunner-Munzel Test ANOVA-type statistics | N2, VEH, *n* = 84 | | 1.38 ± 0.26 |  |  |  |  |  | |
|  |  |  |  | N2, DOI 1 mM, *n* = 65 | | 0.33 ± 0.10 |  |  |  |  |  |  |
|  |  |  |  | *mod-1(ok103)*, VEH, *n* = 55 | | 1.51 ± 0.29 |  |  |  |  |  |  |
|  |  |  |  | *mod-1(ok103)*, DOI 1 mM, *n* = 55 | | 0.67 ± 0.16 |  |  |  |  |  |  |
|  |  |  | Main effect of Strain |  | |  | *F*(1,217.06) = 9.11 | **0.003** | **/† | Cliff’s Δ | 0.16 | |
|  |  |  | Main effect of DOI |  | |  | *F*(1,217.06) = 47.14 | **6.84e^-11^** | ***/††† | Cliff’s Δ | -0.44 | |
|  |  |  | Interaction, Strain × DOI |  | |  | *F*(1,217.06) = 1.50 | 0.222 |  |  |  | |

**Symbols.** *n* indicates that the unit of replication was single worms; *n*^*^ indicates the unit of replication was a cohort of 10-15 worms. The notation e^-x^ represents 10^-x^. Significant *p* values (*p* < 0.05) are in bold text. *Symbols*: **, *p* < 0.01; ***, *p* < 0.001; †, *p* < 0.05 after Bonferroni correction; †††, *p* < 0.001 after Bonferroni correction.
