## Supplemental Table 2 for "Behavioral and genetic analysis of the effects of the psychedelic 2,5-dimethoxy-4-iodoamphetamine (DOI) in *C. elegans*"

| **Metric** | **Fig.** | **Unit of replication** | **N1** | **N2** | **Minimum detectable effect size** | **Empirical pooled std. dev.** | **Minimum detectable difference between means** |
| --- | --- | --- | --- | --- | --- | --- | --- |
| Crawl speed | 2A | Cohort | 31 | 8 | 1.14 | 34.4 | 39.2 μm/sec |
| Swim freq. | 2B | Worm | 88 | 98 | 0.414 | 22.4 | 9.3 waves/min |
| Egg laying | 3 | Cohort | 32 | 12 | 0.970 | 1.22 | 1.2 eggs/worm |
