## Supplementary figures and images for "Behavioral and genetic analysis of the effects of the psychedelic 2,5-dimethoxy-4-iodoamphetamine (DOI) in *C. elegans*"

### Supplemental figure 1

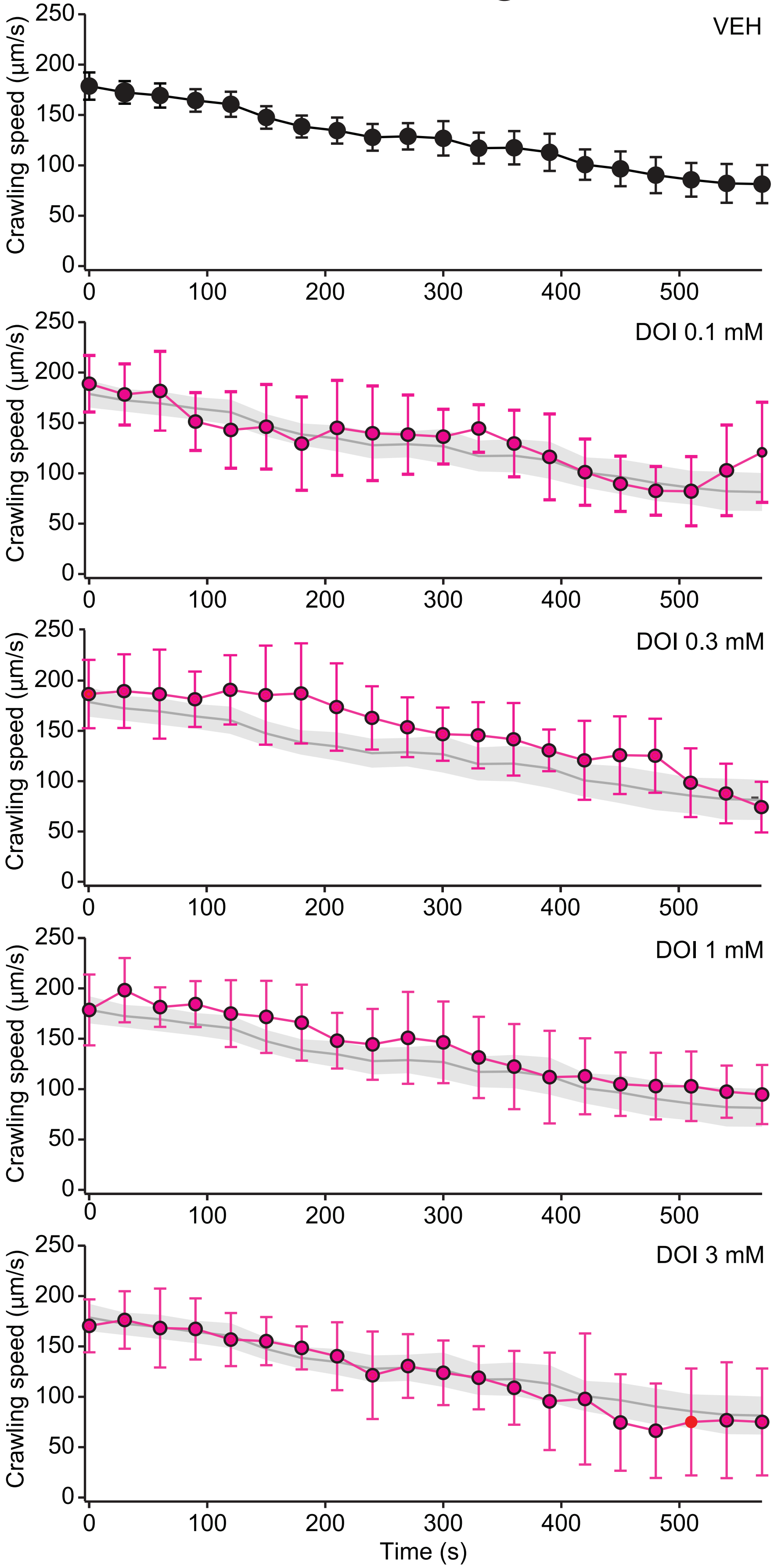

### Supplemental figure 2

#### S2 Fig

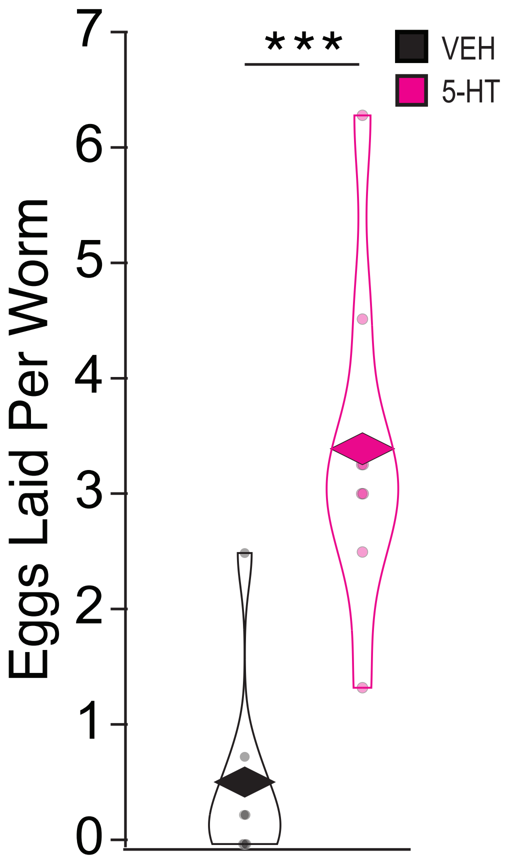

### Supplemental figure 3

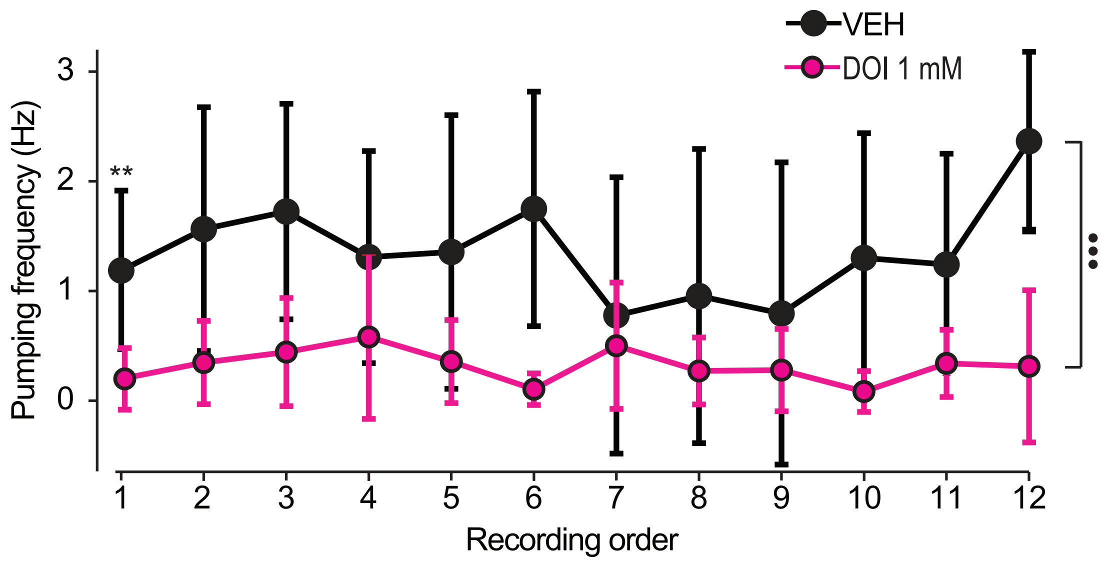
